# Nonlinear sound-sheet microscopy: imaging opaque organs at the capillary and cellular scale

**DOI:** 10.1101/2024.07.31.605825

**Authors:** Baptiste Heiles, Flora Nelissen, Dion Terwiel, Byung Min Park, Eleonora Munoz Ibarra, Agisilaos Matalliotakis, Rick Waasdorp, Tarannum Ara, Pierina Barturen-Larrea, Mengtong Duan, Mikhail G. Shapiro, Valeria Gazzola, David Maresca

**Affiliations:** Department of Imaging Physics, Delft University of Technology, 2628 CJ Delft, the Netherlands; Netherlands Institute for Neuroscience, 1105BA Amsterdam, the Netherlands; Division of Chemistry and Chemical Engineering, California Institute of Technology; Pasadena, CA 91125, USA; Andrew and Peggy Cherng Department of Medical Engineering, California Institute of Technology, Pasadena, CA 91125, USA; Howard Hughes Medical Institute, Pasadena, CA 91125, USA

## Abstract

Light-sheet fluorescence microscopy has revolutionized biology by visualizing dynamic cellular processes in three dimensions. However, light scattering in thick tissue and photobleaching of fluorescent reporters limit this method to studying thin or translucent specimens. Here we show that non-diffractive ultrasonic beams used in conjunction with a cross-amplitude modulation sequence and nonlinear acoustic reporters enable fast and volumetric imaging of targeted biological functions. We report volumetric imaging of tumor gene expression at the cm^3^ scale using genetically encoded gas vesicles, and localization microscopy of currently uncharted cerebral capillary networks using intravascular microbubble contrast agents. Nonlinear sound-sheet microscopy provides a ∼64x acceleration in imaging speed, ∼35x increase in imaged volume and ∼4x increase in classical imaging resolution compared to the state-of-the-art in biomolecular ultrasound.

## Introduction

The most informative method for observing dynamic cellular processes *in vivo* in 3D uses light sheet microscopy that leverages genetically encoded fluorescent reporters (*1*). Successive advances in light sheet microscopy now enable fast, large-volume, and high-resolution imaging of fluorescently labelled cells in transparent or cleared organisms (*2, 3*). These capabilities have had a tremendous impact on developmental biology by enabling long-term imaging of embryogenesis (*4*–*7*).

A next frontier would be to achieve non-toxic deep tissue imaging with cellular precision in living opaque organisms. Unfortunately, limitations inherent to optical microscopy (penetration depth < 1 mm and phototoxicity)(*8, 9*) prevent large scale imaging in opaque tissue. In addition, fast light sheet imaging (*2*) does not yet reach 1 mm^3^s^-1^ volume rates in living tissue (*10*), which makes dynamic imaging at the mesoscale (*11*) technically challenging.

The introduction of biogenic gas vesicles (GVs) (*12*) as the “green fluorescent protein for ultrasound” provides an alternative to light for large-scale cellular imaging. The fortuitous physics of ultrasound enables centimeters-deep scanning of mammalian tissue, while genetically encoded GVs can interface ultrasound waves with cellular function (*13, 14*). To unlock the potential of GV-based acoustic reporter genes (ARGs) (*15, 16*) and biosensors (*17*), there is a need for ultrasound imaging methods with high information content, resolution, coverage, and translatability. Recently, 4D functional ultrasound neuroimaging (*18, 19*) and 3D ultrasound localization microscopy (*20*) have positioned ultrasound as a tool for basic biology research. However, it remains impossible to visualize cellular function in three dimensions or to detect capillary networks.

Here we introduce nonlinear sound-sheet microscopy (NSSM), a method to image targeted biological functions across cm^3^ of opaque living tissue. NSSM relies on a large field-of-view, high-frequency row-column addressed transducer array (RCA) (*21*–*23*) and the transmission of non-diffracting ultrasound beams to detect cells labelled with GVs (*24*) or vessels labelled with microbubbles (MBs) (*25*). NSSM expands the field of view of biomolecular ultrasound at 15 MHz from ∼3.5 × 64 × 100 λ^3^ to 80 × 80 × 100 λ^3^, λ denoting the ultrasound wavelength, the upper bound 2D imaging speed from 400 Hz to 25.6 kHz, and spatial resolution from 1 × 1 × 3.5 λ ^3^ to 1 × 1 × 0.6 λ^3^. We demonstrate the versatility of NSSM by performing volumetric imaging of gene expression in a cancer model, and nonlinear acoustic sectioning of the living cerebral vasculature down to the capillary scale. Throughout the study, NSSM is compared to linear imaging as reference.

## Concept

In the NSSM paradigm, a sub-aperture of RCA-transducer elements *N*_*ap*_(Fig. 1A) is used to transmit cross-propagating ultrasound plane waves, or X-waves, from two adjacent half-apertures at *α* and -*α* angles (Fig. 1B). This spatially structured ultrasound transmission gives rise to a non-diffractive acoustic pressure field in the XZ plane (*26*) exhibiting a double acoustic pressure along the main-lobe of the beam, and a plane wave acoustic pressure field in the YZ plane (*27*) (Fig. 1C). Acoustic pressure is further modulated along the main lobe of the non-diffractive beam using a cross amplitude modulation (xAM) pulse sequence (*24*) (Fig. 1D), which confines nonlinear scattering to a thin sound sheet with a constant beam width regardless of depth (Fig. 1E). The sound sheet beam extends up to the cross-propagation depth 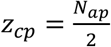 *cotα* which is typically of the order of 100*λ*. 2D images are reconstructed from backscattered ultrasound echoes received on elements of the orthogonal RCA array (Fig. 1B) using a delay-and-sum beamforming algorithm (see Methods). The point spread functions (PSF) of resonant MBs are reported (Fig. 1F) for linear imaging or SSM (i.e. transmission TX1 only) and NSSM (i.e. transmissions TX1-TX2-TX3). We chose MBs as nonlinear point targets in simulation rather than GVs as equations governing their vibration in an ultrasound field are known. XZ-plane SSM and NSSM images showed similar PSFs as both imaging modes operate at the same frequency (Fig. 1G). To scan a volume, sound-sheet transmissions are swept electronically along the two arrays of the RCA-transducer using a sliding aperture of elements (Fig. 1H). A ∼*λ*/2 micro-scanning precision is achieved by interleaving transmissions with and without a silent element at the center of the sub-aperture (Fig. 1H), as the pitch *p* or inter-element spacing of the RCA is approximately equal to *λ*. 3D PSFs are reported in Fig. 1I. Nonlinear imaging is shown to reduce the 3D PSF secondary lobes levels by 12.8dB +/-4.4dB thanks to the confinement of nonlinear MB scattering to the sound sheet plane. In terms of resolution, NSSM delivered an average 3D point spread function of 1*λ* × 0.6*λ* × 0.6*λ* compared to 1*λ* × 0.9*λ* × 0.9*λ* for SSM (Fig. 1J). Additional data supporting Fig. 1 are provided in the Methods section and supplementary Fig. 1.

**Fig. 1.**
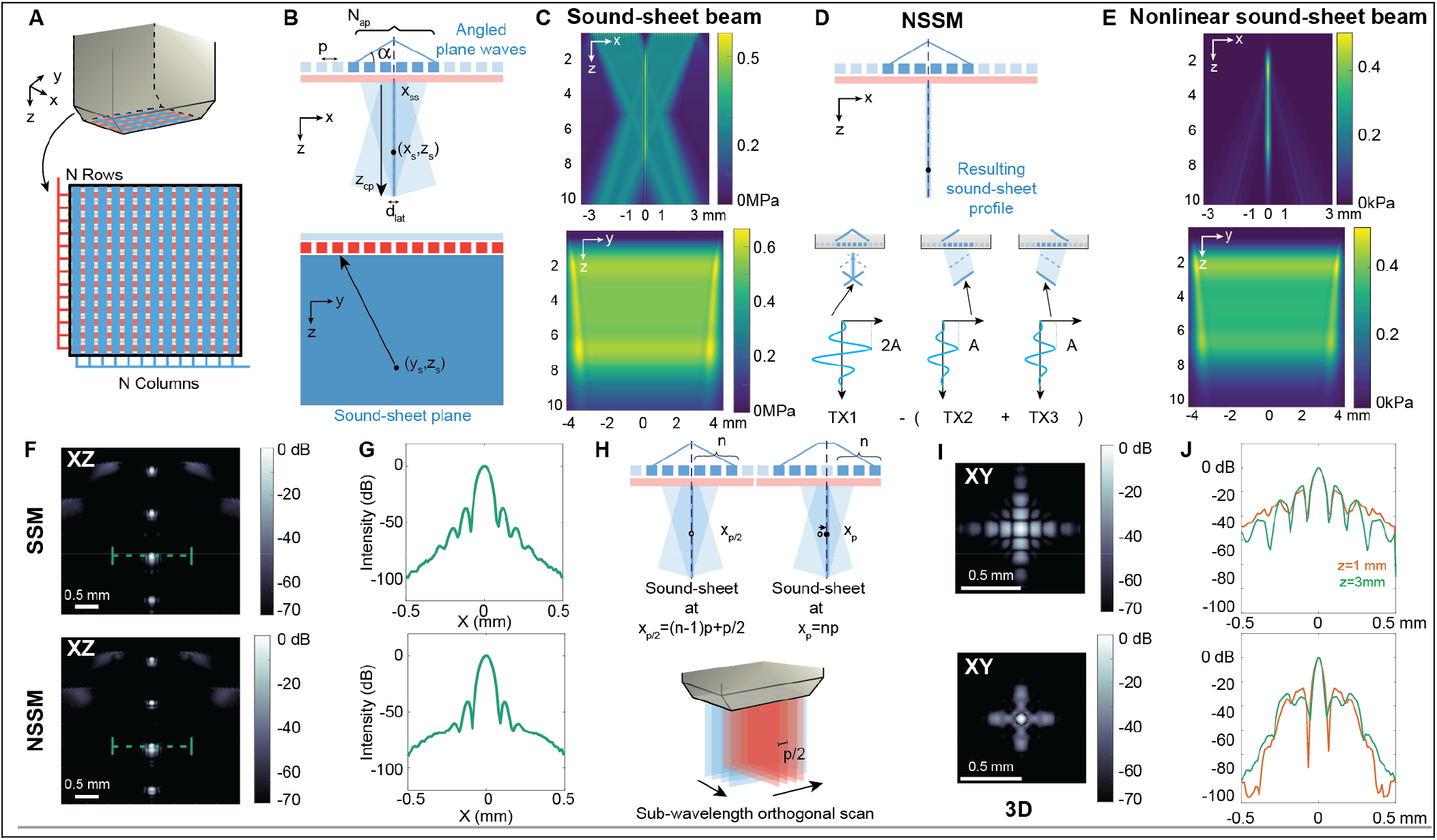
Concept of nonlinear sound-sheet microscopy (NSSM) **(A)** RCA design featuring long and thin transducer elements arranged as rows (red) and columns (blue). **(B)** Top, X-wave transmission with a sub-aperture of column elements. Bottom, reception of backscattered echoes with row elements. **(C)** Simulated X-wave acoustic pressure fields in the XZ and YZ direction. **(D)** Top, spatial confinement nonlinear ultrasound intensity along the main lobe of the X-wave beam using a xAM pulse sequence. Bottom, 3 pulses of the xAM sequence that modulate acoustic pressure *A* along the main lobe of the X-wave beam. **(E)** Simulated residual acoustic pressure field resulting from the xAM sequence in a homogeneous nonlinear water medium. **(F)** Simulated images of four resonant MBs in a water medium. Top, SSM image obtained from transmit event TX1. Bottom, NSSM image obtained from all transmissions of the xAM sequence. **(G)** Top, SSM and bottom, NSSM intensity profiles through a resonant MB. **(H)** Top, electronic sound-sheet micro-scanning along one array of the RCA. Bottom, orthogonal sound-sheet micro-scanning process. **(I)** Top, SSM and bottom, NSSM orthogonally-scanned PSFs of a resonant MB in the XY plane. **(J)** Top, SSM and bottom, NSSM intensity profiles through the center of the 1^st^ and 3^rd^ MB in the X direction.

## Results

Nonlinear ultrasound imaging is key to detect GVs in a tissue context with high specificity (*24*). To test the ability of NSSM to visualize genetic expression in 3D, we first imaged GVs adapted as nonlinear bacterial ARGs (*15, 28*)(Fig. 2). We purified *Anabaena flos aquae* GVs and embedded them in acoustically transparent phantoms at concentrations ranging from optical densities (OD) at 500 nm of 0.5 to 2, thereby mimicking increasing expression levels in cells (*15*) (Fig. 2A). The phantom was scanned with an orthogonal NSSM sequence using a 55*μm* scanning step. SSM detected GV wells at all concentrations (Fig. 2B) and normalized contrast-to-noise ratios (CNR) scaled by steps of 6.9 dB on average from -20.8 dB for OD 0.5 to 0 dB for OD 2. NSSM detected GV wells at all concentrations as well (Fig. 2C) and exhibited a larger, 33 dB dynamic range between the GV well at OD 0.5 and the GV well at OD 2. We observed a lower increase in CNR from OD1.5 to OD2 which may be due to pressure-dependent ultrasound attenuation in a medium containing buckling GVs(*29*). Additional results for linearly and non-linearly scattering purified *Anabaena flos aquae* GVs are reported in Supplementary Fig. 2.

**Fig. 2.**
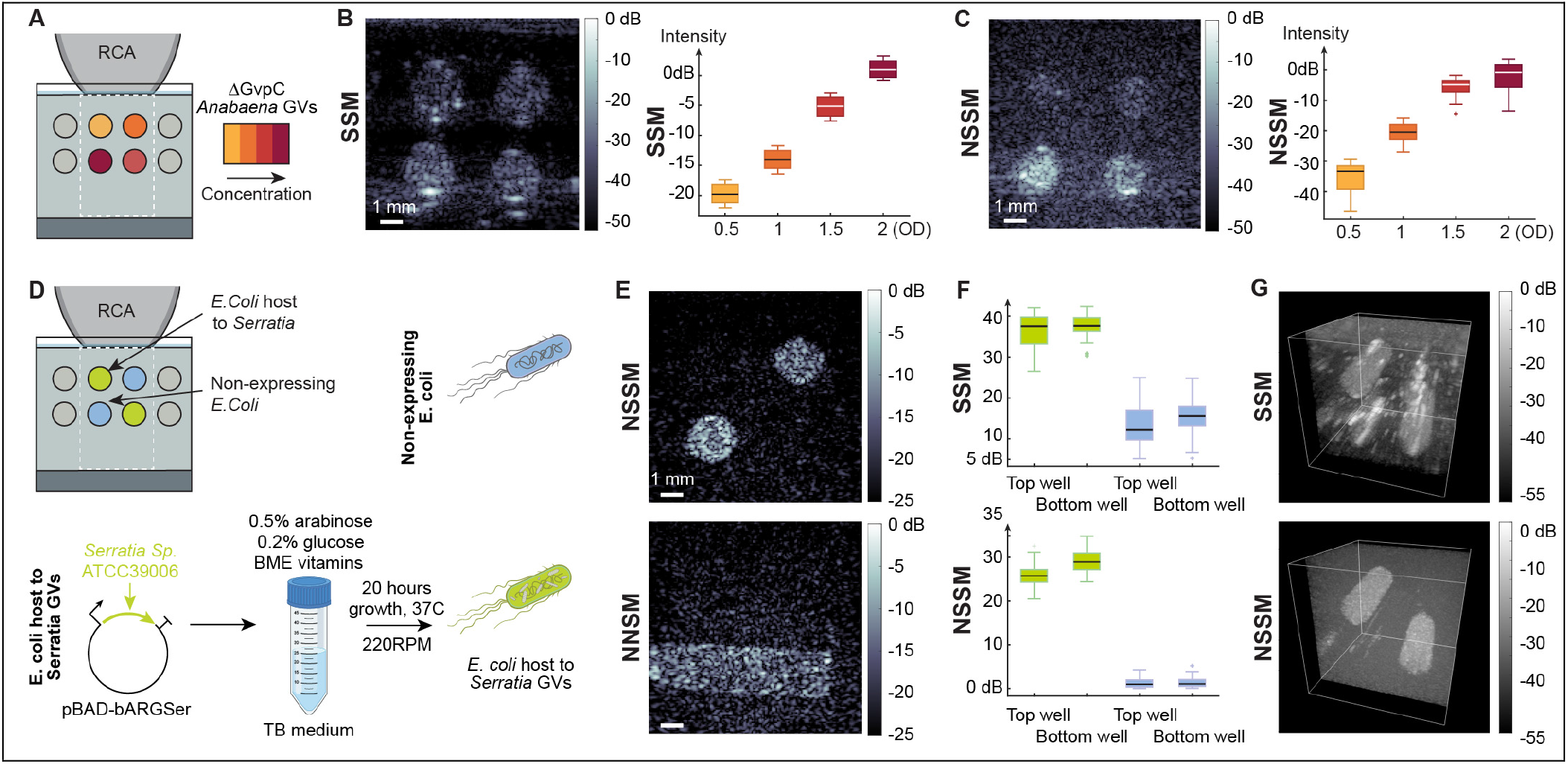
Large volume NSSM of bacterial ARGs **(A)** Experimental setup showing the RCA-transducer in contact with an agar phantom (gray) containing wells filled with increasing concentrations *Anabaena flos aquae* GVs stripped of the shell-stiffening protein GvpC to exhibit nonlinear ultrasound scattering. The RCA field of view is indicated in dashed white lines. **(B)** Left, SSM image for an angle *α=* 19 degrees. Right, contrast-to-noise ratio of each well as a function of the GV optical density. **(C)** Left, corresponding NSSM image. Right, nonlinear contrast-to-noise ratio of each well as a function of the GV optical density. **(D)** Experimental setup with wells containing two strains of *Escherichia coli* bacteria, wild type *E. coli* (blue) and GV-expressing *E. coli*. (green). See protocol (*28*) for bacterial ARG expression. **(E)** Short axis and long axis NSSM images obtained with rows and columns of the RCA respectively. **(F)** SSM and NSSM contrast-to-noise ratios measured out of 108 sound sheet positions across the GV wells. **(G)** 8.8 × 8.8 × 10 mm^3^ volumetric SSM and NSSM images of the phantom.

Next, we tested the ability of NSSM to visualize bacterial ARG expression, which is of particular interest for the field of engineered bacterial biosensors (*30*) and therapeutics (*28, 31*). Two different strains of *E. coli* were used, a control strain and a strain transfected with plasmid pBAD-bARGSer (*28*) leading to intracellular production of *Serratia* GVs, which constitutively produce nonlinear scattering (Fig. 2D). Both strains of bacteria were embedded in agar phantoms and imaged with NSSM (Fig. 2E). Imaging volumes of 8.8 × 8.8 × 10 mm^3^ were reconstructed from 108 sound sheets scanning positions of the RCA-transducer (Fig. 2G). As expected, the control strain did not show any nonlinear contrast, whereas bacteria expressing *Serratia* GVs were detected both in 2D and in 3D (Fig. 2E-G). NSSM detected nonlinear bacterial ARGs with a CNR of 27 dB.

To go further, we investigated the ability of NSSM to image mammalian ARGs (mARGs) in a mouse model of cancer. Orthotopic tumors were induced bilaterally in the mammary fat pads of female immuno-compromised mice by injecting cancer cells engineered to produce nonlinearly scattering GVs (*28*) (Fig. 3A). *In vivo* mARG expression was induced via doxycycline injections every day until day 4 or day 8. Tumors were imaged at day 4 after induction in 3 mice and at day 8 after induction in 2 mice.

**Fig. 3.**
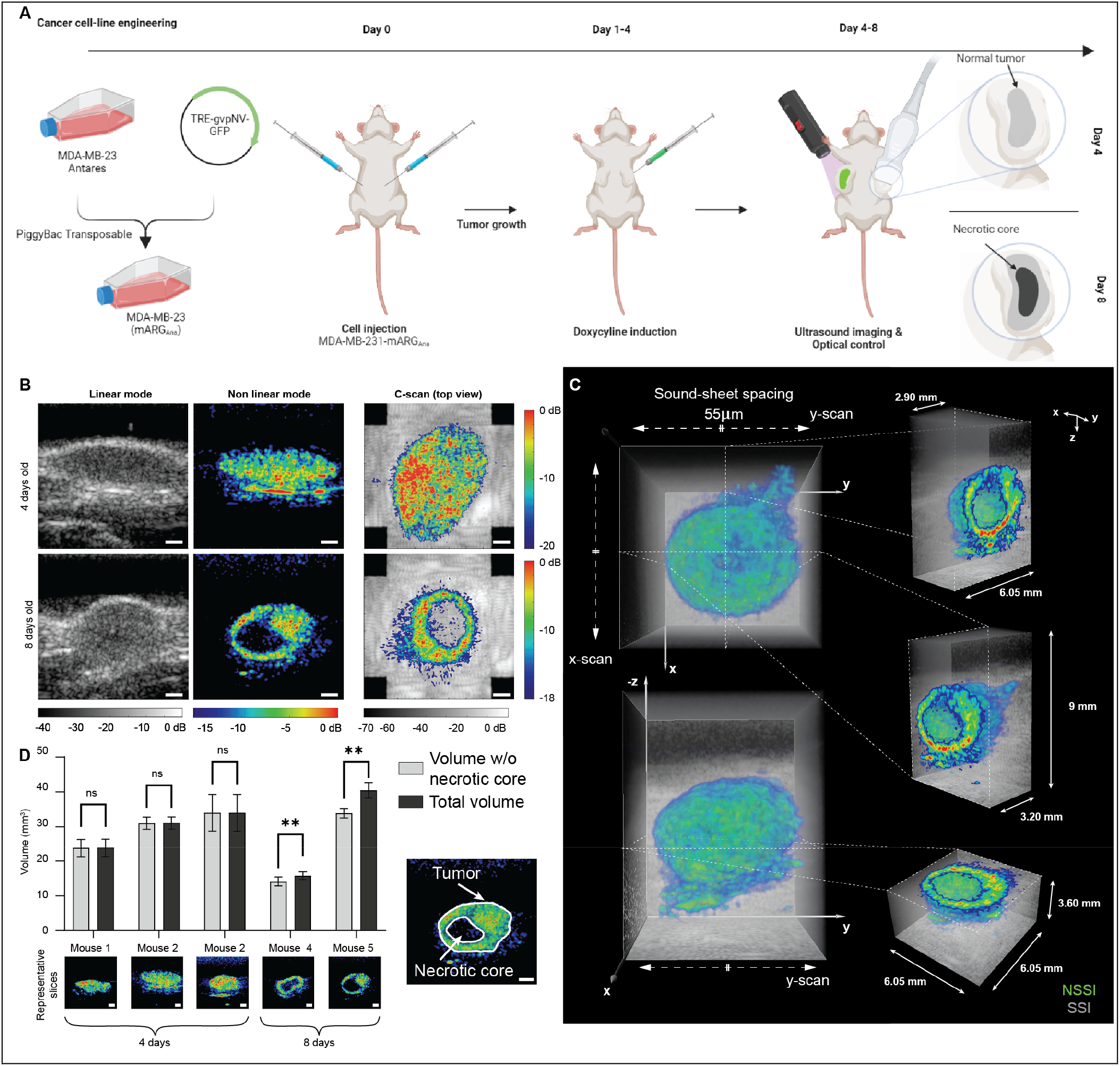
Longitudinal NSSM of tumor gene expression **(A)** Protocol for ultrasound imaging of mARG expression in orthopic tumors (see Methods). **(B)** Left, SSM images of 4- and 8-days old tumors. Middle, NSSM images revealing mARG expression in tumors. Right, XY image of an orthogonal 3D NSSM scan overlaid on linear imaging data. Scalebars, 1 mm. **(C)** Representative 3D NSSM of an 8-days old tumor overlaid on the grayscale SSM volume. **(D)** Results of an automatic segmentation of the tumor and necrotic core for N=5 mice. Light gray plots the tumor volume without the necrotic core, dark gray plots the tumor volume including the necrotic core (multiple paired t-test, \*\**P* < 0.01). Below, representative 2D NSSM of the tumors for each mouse. Scalebars, 1 mm. Right, automatic segmentation result at day 8. Scalebar, 1 mm.

While SSM images revealed anatomical structures, including tumor masses, NSSM successfully revealed spatial patterns of mARG-expression within these tumor masses (Fig. 3B). At day 8 after induction, NSSM revealed the necrotic core of breast tumors via the absence of gene expression. The high specificity of NSSM (*24*), which is robust to nonlinear wave propagation artifacts, was key for this experimental observation. Volumetric imaging enabled to display cross sectional views of mARG expression in the XY plane, referred to as a C-scan (Fig. 3B, right column). Tumors were clearly detected at both stages but necrotic cores were only visible at day 8 after induction.

Volumetric NSSM fused with anatomical SSM imaging is reported in Fig. 3C. The total volume scanned extends across 8.8 × 8.8 × 9 *mm*^3^ and was acquired with a with scanning step of 55 *μm* along each array of the RCA-transducer. Fig. 3C displays cross sectional views of gene expression in XZ, YZ and XY planes, illustrating 3D navigation capabilities of NSSM. We quantified tumor and necrotic core volumes using a custom automatic segmentation pipeline (Fig. 3D). At day 4 after induction, volumes measured with and without necrotic cores were similar whereas at day 8 after induction, volumes measured with and without necrotic cores showed a statistically significant difference. A representative automatic segmentation of tumor gene expression contours is provided in Fig. 3D. In this deep tissue imaging example, given the static nature of gene expression at our imaging speed, 2D NSSM was operated at a 930 frames/s whereas orthogonally scanned volumetric NSSM was operated at a 4 volumes/s. In theory though, 3D NSSM could reach 94 volumes/s for this set of imaging parameters. Interestingly, the quantification of *in vivo* tumor volumes would have been impossible based on anatomical ultrasound imaging alone, which highlights the potential of this imaging method.

Alongside genetically encoded GVs, synthetic lipid-shelled MBs are another class of ultrasound contrast agents used as vascular reporters. MBs exhibit amplitude-dependent ultrasound scattering (*32*), which makes them detectable with amplitude modulation pulse sequences as well (*33*). To test the capacity of NSSM to visualize MBs circulating in blood vessels, we performed high-speed nonlinear Doppler imaging of the rat brain vasculature (Fig. 4).

**Fig. 4.**
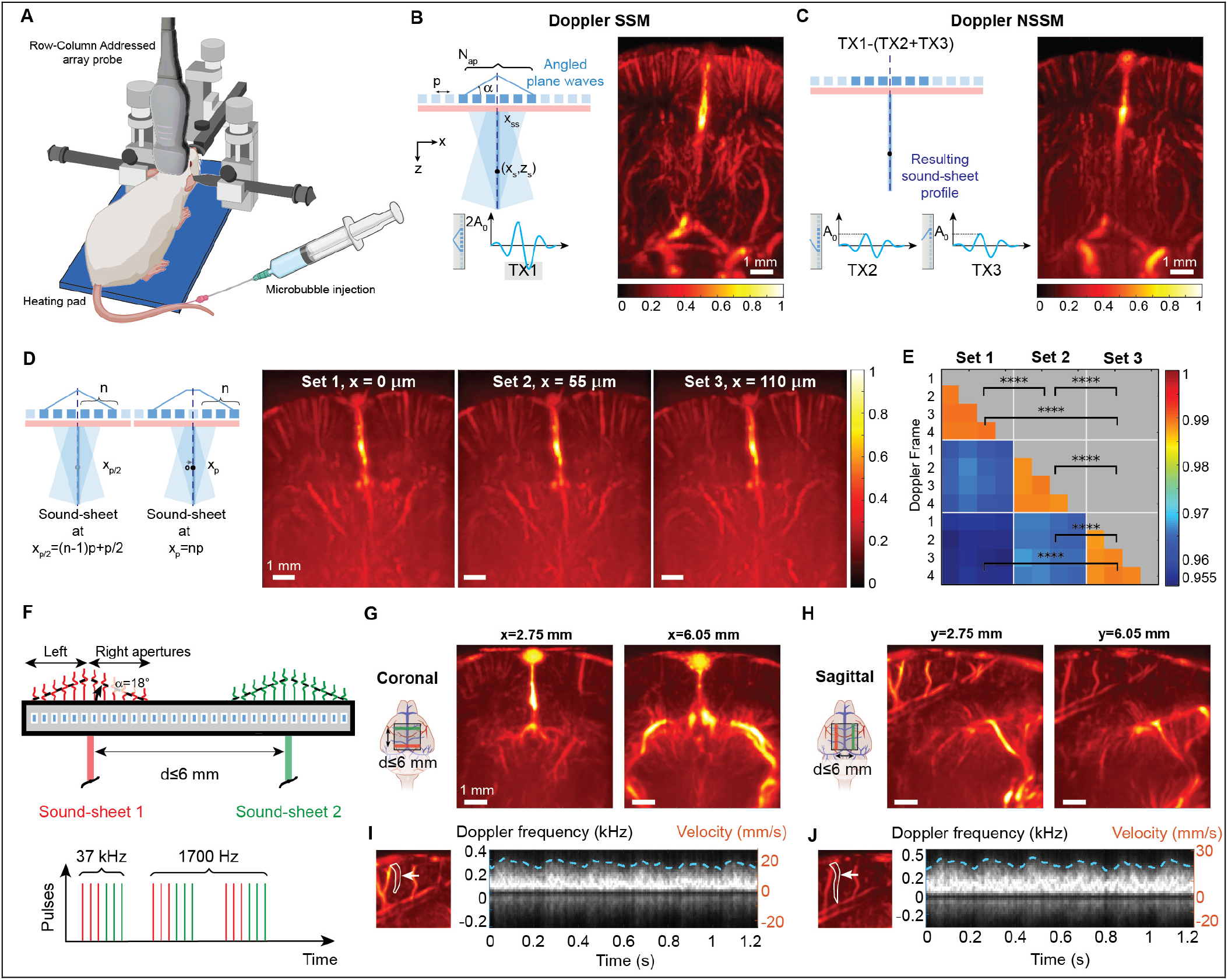
High-speed multi-view NSSM of the rat brain vasculature **(A)** Experimental setup. **(B)** Coronal section of the rat brain vasculature acquired with 4.4 kHz Doppler SSM after MB injection. **(C)** Same brain section acquired with 4.4 kHz Doppler NSSM after MB injection. **(D)** Doppler NSSM of three adjacent coronal brain sections with a 55*μm* micro-scanning step. **(E)** Structure Similarity Index Matrix calculated using 4 Doppler NSSM acquisition in each of the three adjacent planes (\*\*\**P* < 0.001, **** *P* < 0.0001). **(F)** Multi-view NSSM imaging principle using two sub-aperture of elements of the RCA-transducer. With our RCA probe, imaging planes can be spaced up to 6 mm, and imaging speed was set to 1.7 kHz per plane. **(G)** High-speed multi-view Doppler NSSM of two coronal planes. **(H)** High-speed multi-view Doppler NSSM of two symmetric sagittal planes. **(I)** Doppler spectrum revealing peak cerebral blood flow and pulsatility of a cortical arteriole of the left hemisphere. **(J)** Doppler spectrum revealing peak cerebral blood flow and pulsatility of a cortical arteriole of the right hemisphere.

MBs tuned for high-frequency ultrasound were administered via tail vein injections in anesthetized head-fixed rats (Fig. 4A). As reference, linear Doppler SSM images were acquired at a 4.4 kHz framerate (Fig. 4B) and generated results similar to ultrafast Doppler images of the rat brain (*34*). Doppler NSSM images (Fig. 4C) were obtained using amplitude modulated data and a high-pass filter to remove residual static echoes (see Methods). Note that nonlinear Doppler results did not rely on any singular value decomposition (SVD) filter (*35*). Because nonlinear Doppler processing spatially confines image data to a 100 µm x 9.6 mm x 8.8 mm thin sound sheet plane, we detected less vessels in Fig. 4C than in Fig. 4B, which projects in one image echoes arising from the oblique paths of each plane waves (see Fig 1C). As a result, the cortical surface was clearly delineated in Doppler NSSM (Fig. 4C) whereas vascular signals projected from the oblique paths of each plane waves are visible above the cortex in Fig. 4B. Supplementary Fig. 3 shows that Doppler NSSM is more sensitive to slow blood flows than Doppler SSM based on SVD filtering, in line with previous observations (*36*).

Next, we performed ultrasound sectioning the rat brain vasculature with sub-wavelength scanning steps of 55 *μm* (Fig. 4D), acquiring four consecutive Doppler acquisitions per plane. We quantitatively assessed vascular changes in these adjacent planes by computing structural similarity index matrix values for each Doppler acquisition (Fig. 4E). Intra-sets, the structural similarity index averaged to 0.99 indicating that vascular images were nearly identical across cardiac cycles. Inter-sets, the index dropped to 0.96 confirming that we observed two separate vascular planes.

Inspired by multi-slice tomography techniques, we tested the ability of Doppler NSSM to capture multiple views of the brain simultaneously. To do so, we interleaved pulse sequence transmission using two sub-apertures of elements of the array (Fig. 4F and supplementary Fig. 4). In this configuration, we set our imaging rate to 1.7 kHz in both sound-sheet planes. To demonstrate the versatility of this approach, we imaged two coronal vascular planes separated by 3.3 mm using the first array of the RCA-transducer (Fig. 4G). Similarly, we imaged two sagittal vascular planes separated by 3.3 mm using the second array of the RCA-transducer, revealing symmetric vascular planes in each hemisphere (Fig. 4H). Last, we processed SSM Doppler spectrograms in each sagittal plane to show that acquisitions were continuous and co-registered in time (Fig. 4I-J). The Doppler-derived heartrates were equal to 298 bpm and 295 bpm in each brain hemisphere, respectively.

In 2015, vascular ultrasound was redefined by the introduction of ultrasound localization microscopy (ULM), a super-resolution method that can map the *in vivo* microvasculature with a ∼*λ*/8 resolution (*37*). A drawback of current ULM processing is that SVD-based filters used to isolate MB echoes are unable to detect slowest blood flow velocities occurring in capillary beds (*36, 38*). As a consequence, ULM cannot visualize capillaries so far while it is the vastest vascular territory of living organisms (*39*). Since the NSSM detection of MBs relies on nonlinear MB scattering rather than MB motion, we hypothesized that the combination of NSSM and ULM could potentially reveal capillary beds *in vivo*.

We investigated nonlinear sound-sheet localization microscopy (NSSLM) of the cerebral capillary vasculature in Fig. 5. Craniotomized rat brains perfused with MBs were imaged at 1 kHz using NSSM in a coronal mid-brain plane over the course of 105 seconds, leading to the acquisition of 10^5^ frames (Fig. 5A). Two post-processing pipelines were used to generate state-of-the-art ULM images and NSSLM images. Briefly, NSSLM processing consisted in filtering MB echoes with the amplitude modulation step in NSSM, followed by velocity filtering to isolate MB in several velocity bands [0-3], [3-15], [15-150] mm/s. The position of individual MB echoes was estimated with a radial symmetry algorithm and trajectories were reconstructed using Kuhn-Munkres pairing (*40*). Fig. 5B displays SSM Doppler and NSSM Doppler images filtered in the capillary flow velocity band (0.5 to 3 mm.s^-1^) (*41, 42*), showing that NSSM retrieves vascular signals in cortical and hippocampal regions of the rat brain with a good SNR, whereas the SSM images is mostly filled with diffuse vascular noise. In particular, low velocities located at the wall of the sinus vein are visible in Doppler NSSM. Time-series of NSSM frames filtered in the capillary velocity band (Fig. 5C) show the dynamic of slow flowing MBs captured with NSSM, which constitutes the basis for NSSLM post-processing. Individual microbubbles that are quasi-static are indicated with white arrows. Over the course of 75 ms, several MBs progress by less than half a wavelength (57 *μm*), indicating that their velocities is below 0.8 mm/s which falls in the range of capillary flow velocities.

**Fig. 5.**
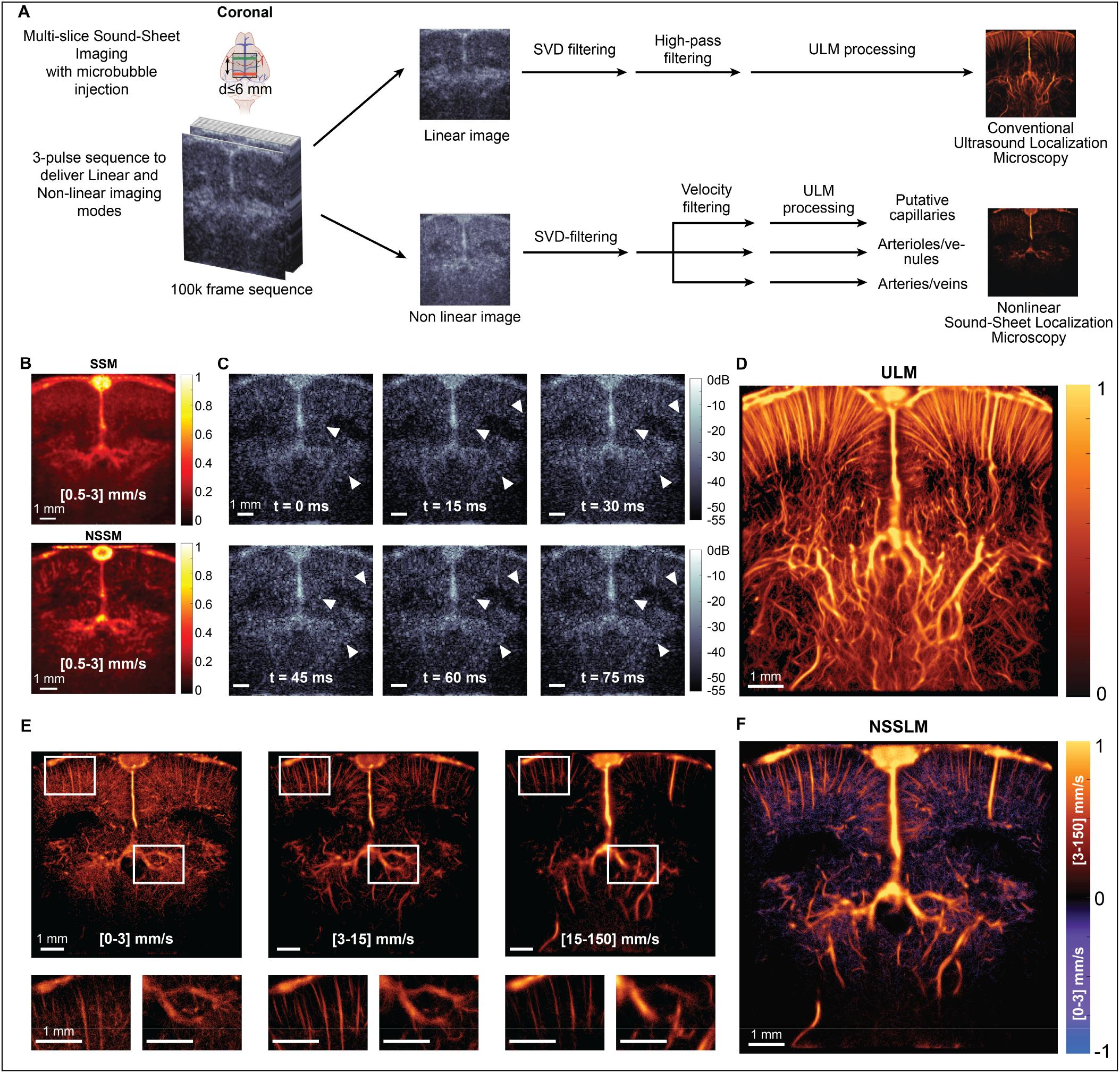
Nonlinear sound-sheet localization microscopy of rat brain capillaries **(A)** SSLM and NSSLM image processing pipelines. **(B)** Vascular information captured by Doppler SSM and Doppler NSSLM in the capillary blood flow velocity band (0.5-3mm.s^-1^). **(C)** NSSM frames of the NSSLM of the NSSLM sequence revealing slow MB flow in the rat brain. **(D)** ULM image generated with a state-of-the-art ULM processing pipeline. **(E)** NSSLM in the capillary flow velocity band (0.5-3 mm.s^-1^), arteriole and venule flow velocity band (3-15 mm.s^-1^) and artery and vein velocity band (15-150 mm.s^-1^). **(F)** Composite NSSLM image display all cerebral blood flow velocity bands.

As a reference, we processed a state-of-the-art ULM density map using transmission TX1 of the NSSM sequence (Fig. 5D). In comparison, NSSLM (Fig. 5E-F) enabled mapping of capillary beds segmented with a 0-3 mm.s^-1^ flow velocity band (Fig. 5E, left panel), arterioles and venules segmented with a 3-15 mm.s^-1^ flow velocity band (Fig. 5E, middle panel) and arteries and veins segmented with a 15-150 mm.s^-1^ flow velocity band (Fig. 5E, right panel). A composite NSSLM image showing all vascular compartments is presented in Fig. 5F. Vascular structures displayed in the NSSLM image appear clearly denser from structures detected with SSLM, as expected from capillary beds (*39*). The results for the second plane exhibit similar characteristics and are presented in supplementary Fig. 6 and 7.

## Discussion

We report NSSM, a fast and volumetric imaging method capable of visualizing targeted biological processes at the organ scale. NSSM introduces nonlinear sound-sheet beams in the field of ultrasound imaging and is capable of detecting two major classes of ultrasound contrast agents, genetically encoded GVs and intravascular MBs. Thanks to its all-acoustic nature, NSSM circumvents certain limitations of light-sheet or multi-photon microscopy (*3, 43, 44*) such as phototoxicity, photobleaching of fluorescent reporters and complex single-objective microscope designs based on oblique illumination.

In 2D, NSSM achieved framerates of 4.4 kHz in thin sound-sheets of 0.1 × 8.8 × 12.9 mm^3^, or 1 × 89 × 130 λ^3^ (see Fig. 4D). We show that sound-sheet beams can be arbitrarily positioned within the 8.8 ×x 8.8 mm^2^ active aperture of a 15 MHz RCA-transducer and swept electronically along each orthogonal array with a sub-wavelength precision of 55 µm (∼*λ*/2).

NSSM can be further tuned for speed or spatial coverage. Inspired by multi-slice imaging of the brain (*45*), we report fast multi-view NSSM at a framerate of 1.7 kHz, enabling Doppler imaging of the rat brain. Interestingly, symmetric planes in each brain hemisphere can be observed simultaneously and enable studies of lateralized brain function. The 64x increase in 2D imaging speed compared to xAM imaging enabled localization microscopy of the cerebral capillary vasculature as shown in Fig. 5. Here, the combination of kHz imaging with specific MB detection regardless of MB velocity was fundamental to chart capillary territory and opens the way for imaging of microvascular activity deep in intact tissues. In this first NSSLM demonstration, we acquired 2D super-resolution vascular images in parallel coronal planes spaced by several millimeters. Future work will investigate the feasibility of 3D NSSLM of the capillary vasculature using fast electronic sweeping of sound-sheet beams along adjacent planes spaced by half a wavelength.

Orthogonal NSSM microscanning enables volumetric imaging of genetically encoded bacterial (Fig. 2) and mammalian ARGs (Fig. 3). In particular, 3D NSSM enabled longitudinal molecular ultrasound imaging of gene expression in a tumor model. The imaging method creates the possibility for *in vivo* tumor volume quantification in mammalian tissue. The high specificity of cross amplitude modulation was critical to reveal the necrotic core of tumors, as classical pulse sequences would underestimate it due to nonlinear wave propagation artifacts.

The NSSM paradigm will complement the capabilities of current 4D ultrasound imaging methods such as orthogonal plane wave imaging (*22*), synthetic aperture imaging (*21*) or xDoppler (*46*) that provide tissue information at the mesoscale but do not enable molecular or cellular imaging. Compared to 2D ultrafast ultrasound imaging, NSSM provides an improved resolution in elevation which can be as low as 0.4λ (*24*) versus *≥* 3λ (*47*) for 1D transducer arrays. Compared to orthogonal plane wave imaging, the lateral resolution is improved due to the thinner sound-sheet beam in NSSM (see supplementary Fig. 1).

A first limitation of NSSM is that the requirement for symmetric half-apertures to generate sound-sheet beams prevents imaging on the edge of the RCA-transducer. However, orthogonal NSSM microscanning enables to retrieve part of this missing volume. In the end, the surface area lost in the corners of the active surface of the RCA-transducer represents 11% (77.44 *cm*^2^ to 69.24 *cm*^2^). In addition, wider RCA-transducers made of 256 elements could be operated with our current hardware and would automatically increase the field of view.

A second limitation lies in the volumetric imaging rate achievable with SSM which lies in the hundreds of Hz and not kHz. This volume rate remains comparable to other high quality RCA volumetric imaging methods, such as functional OPW neuroimaging (*48*) or synthetic aperture B-mode imaging (*49*).

A third limitation of our study is that brain imaging was performed in craniotomized animals. The transcranial potential of NSSM has not been explored. Meanwhile, NSSM will be compatible with chronic acoustic windows (*50*).

Independently from the method itself, NSSM would also benefit from the development of monodisperse MBs tuned for high frequency ultrasound as it would improve CNR and allow us to reduce MB doses administered (*51*).

The performance of NSSM is also bound by the collapse pressure of ARGs. A tradeoff exists between imaging depth that requires high acoustic pressures, and near field detection of ARGs that would be destructed if high acoustic pressures are used.

To conclude, the combination of latest and future generation acoustic probes (*52*) with NSSM carries a wave of opportunities for deep tissue imaging of dynamic biological processes. NSSM offers an unprecedently high spatiotemporal resolution and coverage to explore living opaque organs across scales.

## Contribution statement

BH and DM conceived the project. BH developed the NSSM sequence, k-Wave simulation, acquired experimental data and developed processing scripts. DT, BMP, EMI, TA optimized and sustained gas-vesicle production. BH performed animal surgeries with significant support from FN, EMI. AM developed INCS simulations. RW contributed to experimental data acquisition and discussions on beamforming optimization. PBL, MD provided cancer models and performed *in vivo* experiments with BH. VG contributed brain research resources. MGS and VG commented the manuscript. BH and DM wrote, reviewed, and edited the manuscript.

## Data and code availability statement

Raw data as well as ultrasound acquisition and processing scripts used to generate key figures will be posted to a publicly accessible GitHub repository at the time of manuscript publication.

## Methods

### Generation of non-diffractive ultrasound beams

RCA rows or columns were used to transmit simultaneous cross-propagating plane waves from two contiguous half-apertures 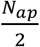 at angles *α* and -*α* (Fig. 1B). The two transmitted plane waves interfere along a 2D plane, referred to as the sound-sheet plane. During cross-propagation, an X-wave with a double amplitude is generated and propagates with a supersonic velocity 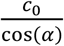 until the cross-propagation depth 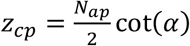. Ultrasound echoes received by the array orthogonal to the transmitting array are processed to generate an image. Image reconstruction relies on a delay-and-sum beamforming algorithm with the assumption that ultrasound backscattering only arises from the sound-sheet plane. For beamforming delay laws, see the image reconstruction paragraph and reference (*25*).

### Sub-wavelength micro-scanning and 3D ultrasound imaging

Non-diffractive beams can be generated two ways (see Fig. 1H). First, using two contiguous sub-apertures of RCA-transducer elements. This focuses the sound-sheet plane at a position 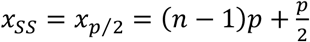, with *n* the number of elements and p the pitch or inter-element spacing of the RCA-transducer. Second, using two sub-apertures of RCA-transducer elements separated by an inactive element. This focuses the cross-propagation plane at a position *x*_*SS*_ = *x*_*p*_ = *np*. In practice, these two transmissions generate non-diffractive beams with main-lobes separated by a distance of 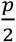. The thickness of the sound sheet was estimated using INCS simulations (*25*), the Full Width at Half Maximum (FWHM) of the main-lobe of the non-diffractive beam can be approximated with the following relation:

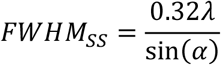

In particular, the FWHM becomes smaller than the wavelength for any angle *α* > 18.66°.

By performing sound-sheet micro-scanning in X and Y, a volume is sampled. Note that sound-sheet micro-scanning can also be perform along only X or Y. Because sound-sheets are generated using two half-apertures, the field of view *FoV* is reduced compared to active surface of the RCA-transducer: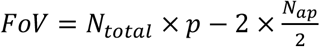. The volume occupied by the main-lobe of a sound sheet beam is therefore: *z*_*cp*_ × *FWHM*_*SS*_ × *w*_*array*_, with *w*_*array*_ = *N*_*total*_*p* the full width of the RCA array. Therefore, the volume of sound-sheet scan along one array is equal to:

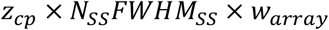

with *N*_*SS*_ the number of sound sheet positions.

By using varying aperture sizes, it is possible to extend the field of view further, but then the propagation depth *z*_*cp*_ is reduced. To keep *z*_*cp*_ constant, one possibility is to decrease angle *α*. Considering a minimum half-aperture for sound-sheet transmission *N*_*red*_*p*, the total field of view covered by a scan in one direction is:

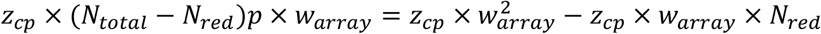

and for a 3D scan along X and Y, this becomes:

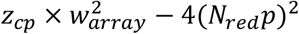

When compared to active surface of the RCA-transducer, this leads to a field of view reduction of:

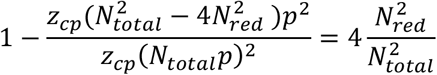

In our study, we used a reduced aperture of as low as 12 elements, leading to a loss 11% loss in image volume.

### Spatial confinement of nonlinear scattering via amplitude modulation

In NSSM, the same half-apertures are used for two additional transmits, each firing independently this time (see events TX2 and TX3 in Fig. 1D). As no cross-propagation is taking place in this case, the acoustic pressure delivered along the sound-sheet plane has a twice lower amplitude than TX1. An amplitude-modulated signal can thus be obtained by subtracting echoes received from TX2 plus TX3 from echoes received from TX1. This operation is done on the radiofrequency data and the residual signal is beamformed and further processed. A 3D NSSM image can also be obtained by performing three xAM pulse transmissions for every sound-sheet position in X and Y.

### Sound-sheet image reconstruction

To reconstruct a sound-sheet image, we make the assumption that backscattered echoes originate only from the sound sheet plane. If the sound sheet plane is oriented with its normal along the 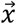 direction, then all scatterers can be assumed to have the same *y* coordinate. The forward delay for a scatterer at a position (*x*_*s*_, *y*_*s*_, *z*_*s*_) can be written as:

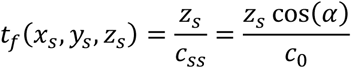

With *c*_*SS*_ the supersonic speed of the X-wave and *c*_0_ the speed of sound in the medium.

The return delay for a scatterer at a position (*x*_*s*_, *y*_*s*_, *z*_*s*_) is calculated for the array orthogonal to the transmitting array to restore focusing and is written as:

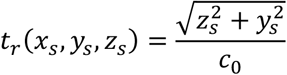

It is important to notice that *t*_f_ only depends on *z*_*s*_, *x*_*s*_ while *t*_r_ only depends on *z*_*s*_, *y*_*s*_.

### Simulation of linear and nonlinear sound-sheet beams

Linear and nonlinear sound-sheet beams reported in Fig. 1C and Fig. 1E were computed with the Iterative Nonlinear Contrast Source (INCS) method (*25*). Briefly, INCS was developed to solve the four-dimensional spatiotemporal Westervelt equation describing nonlinear sound wave propagation. Simulations were conducted within a computational domain *X* × *Y* × *Z* = 6.5 × 13 × 10.5 *mm*^3^. The propagation medium consisted of water and was characterized by a mass density *ρ*_o_ = 1060*kg. m*^-3^ and a speed of sound *c*_0_ = 1482 *m. s*^-1^. The incident beam had a center frequency *f*_*c*_ = 15*MHz*. The simulated RCA array consisted of 64 individual elements, each with length 12.8 *mm*, and a pitch of 100 *μm*. The transmitted pulses were modelled as Gaussian-apodized sine-bursts:

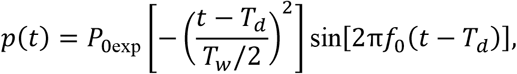

with *T*_*w*_ = 1.5/*f*_*c*_ representing the duration of the Gaussian envelope and *T*_d_ = 3/*f*_*c*_ + Δ_*n*_ a total time delay. The latter consists of a fixed delay for keeping *p*(*t* = 0)∼0, plus a delay per element for the beam steering. Beams are simulated for an angle *α* = 20.7°. The peak acoustic pressure of the elements surface was *P*_0_ = 400 *KPa*. A sampling frequency of 90 MHz was used to discretize the spatiotemporal domain.

### Simulation of NSSM point spread functions

We simulated the response of 4 resonant microbubbles in a homogeneous water medium using the k-Wave simulation toolbox. MBs were positioned at *z* = 1; 2; 3 and 4 *mm* and centered in the middle of the domain. Each MB had a diameter of 1.5*μm*, leading to a resonance frequency of 15.625 *MHz*. In this simulation, the geometry of the transducer consists of 43 elements with a height of 4.3 *mm*, a width of 100 *μm* and a pitch of 100 *μm*, and a bandwidth of 14 to 22 *MHz*. 22 sound-sheet transmissions were simulated with a scanning step of 50 *μm*, an angle *α*=21°, a 4-cycles Gaussian-apodized sine-burst and an acoustic peak pressure of 400 *KPa* at the transducer surface. The radiofrequency data was beamformed using the delay-and-sum algorithm described above. For 3D imaging, we permuted simulation data to emulate data of the orthogonal array as our problem is symmetric.

### Ultrasound imaging of a wire phantom

A 15 MHz RCA probe with 80+80 elements (Verasonics, Kirkland, WA, USA) with a pitch or inter-element spacing of 110 µm was placed over a 3D wire phantom (model 055A from CIRS, VA, USA). The wires were oriented at an angle from the orientation of the rows and columns elements. SSM images were obtained using a 14° angle per array (2 transmits per voxel result) and a combination of [7°, 15°, 20°] angles (6 transmits per voxel result). As reference, an orthogonal plane wave (OPW) acquisition with an angle of -18 degrees was used (2 transmits per voxel result) and 2×32 transmits ranging from -21° to 21° (64 transmits per voxel result). Imaging results are reported in supplementary Fig. 1. C-scans, i.e. images in the XY plane, reveal wires in diagonal. For the SSM case, micro-scanning was performed with a 55 *μm* step. In-plane images, i.e. XZ scans, reveal a single wire cross section. Waveforms transmitted in both case were 13.6 MHz, 0.5 cycles sin-bursts.

### Ultrasound imaging of GV phantoms

**Fig. 2A:** *Anabaena flos aquae* GVs were cultured and transferred to sterile separating funnels. Buoyant cells were separated from the growth media through natural flotation, and GVs were harvested after 48h of hypertonic lysis. A cycle of centrifugation and resuspension allows to purify the GVs further. A stock of *Anabaena flos aquae* GVs was stripped of their GvpC protein layer with a 6-M urea solution to obtain nonlinearly scattering GVs (ΔGvpC GVs). A 2% agar phantom comprising 2 mm in diameter wells was casted using custom-printed molds and imprints. For each of the 4 wells, the concentration is increased starting with a concentration measured optically at OD0.5, OD1, OD1.5 and OD2. Images were obtained with the 15 MHz RCA-transducer (Verasonics, Kirkland, WA, USA). Waveforms transmitted were 13.6 MHz, 0.5 cycles sin-bursts. The X-wave angle *α* was set to 19°.

**Fig. 2D:** *Escherichia coli* bacteria were grown for 20 hours at 37C, 220 RPM flask shaking in a terrific broth medium with 0.5% arabinose, 0.2% glucose and BME vitamins. In one case, plasmid pBAD-bARGSer was transferred to the *E. coli* strain (courtesy of Shapiro Lab, derived from plasmid Addgene #192473) via electroporation. These cells produced *Serratia* GVs that are constitutively nonlinear scatterers. Both bacterial populations were then transferred to the phantom.

### Ultrasound imaging of a cancer model

All *in vivo* experiments were performed under protocol 1761, approved by the Institutional Animal Care and Use of Committee of the California Institute of Technology. Animals were housed in a facility maintained at 71–75 °F and 30–70% humidity, with a lighting cycle of 13 hours on and 11 hours off (light cycle 6:00–19:00). Tumor xenograft experiments were conducted in NSG mice aged 12 weeks and 6 days (Jackson Laboratory). As we relied on an orthotopic model of breast cancer, all mice were female. MDA-MB-231-mARGAna cells were grown in T225 flasks in DMEM supplemented with 10% TET-free FBS and penicillin–streptomycin until confluency as described above. Cells were harvested by trypsinization with 6 ml of trypsin/EDTA for 6 minutes and quenched with fresh media. Cells were washed once in DMEM without antibiotics or FBS before pelleting by centrifugation at 300×g. Cell pellets were resuspended in a 1:1 mixture of ice-cold Matrigel (HC, GFR) (Corning, 354263) and PBS (Ca2+, Mg2+-free) at 30 million cells per milliliter. Then, 50-µl Matrigel suspensions were injected bilaterally into the 4th mammary fat pads at 1.5 million cells per tumor via subcutaneous injection. Twelve hours after tumor injection and every 12 hours thereafter (except the mornings of ultrasound imaging sessions), test mice were intraperitoneally injected with 150 µl of saline containing 150 µg of doxycycline for induction of GV expression.

The 15 MHz RCA probe was operated to transmit at 110 sound-sheets positions, with angles ranging from 15° to 21° with a 1 degree step per sound-sheet position. All 3D renderings were generated using the Avizo rendering software (ThermoFisher ®). To segment and measure the tumor and hypoxic core, the data was first 3D-Gaussian filtered (standard deviation *σ* = 0.6) and interpolated twice in each direction. The data was then normalized, and log-compressed and tissue attenuation was taken into account with an average tissue attenuation factor of 0.54*d B*/*MHz*/*cm*. Attenuation was further corrected to aim for a uniform noise contrast value through depth of the image, an additional custom Gaussian filtered was added pre-binarization to yield better results. The open volume result was then closed and measured using the regionprop function in Matlab. This was done for each of the volumes per acquisition obtained in the two directions of scanning and in the 3D compounded volume. With the approach, we calculated an average value for the volume of the tumor and hypoxic core.

### Ultrasound Doppler imaging

The RCA probe was used to image the vascular function of a rat brain (Sprague Dawley, female, 280g). All experiments were performed under CCD license number AVD8010020209725 at the Koninklijke Nederlandse Akademie van Wetenschappen with Study Dossier number 213601. Immediately after isoflurane induction, carprofen, and butorphanol were delivered subcutaneously. The animal was prepared (shaved, disinfected, placed in earbars) and surgery began no sooner than 20 minutes after injections. A catheter was placed in the tail vein. Heparin was injected to prevent blood clots forming in the catheter. 14×14 mm craniotomies were performed. Carprofen and butorphanol were delivered subcutaneously during surgery (respectively 5mg/kg and 2mg/kg). To prevent cerebral edema, dexamethasone was given subcutaneously with a dosage of 2.5mg/kg.

After craniotomy, the RCA probe was positioned over the cranial window of anesthetized animal. Sound-sheet were transmitted with an 18° angle and 38 elements were used for each half apertures. The transmitted pulses were 2-cycle sine-bursts centered at 15.6 MHz. For Fig. 4c, 2100 frames were acquired with a frame rate of 440Hz. For Fig. 4D, 2100 frames were acquired with a frame rate of 300Hz, leading to a 7s long acquisition. For Fig. 4G and 4H, 2100 frames were acquired with a frame rate of 1700Hz, leading to a 1.24s long acquisition. The average heart rate throughout the experiment is 419 bpm, leading to a cardiac cycle of 143 ms which is short enough to be captured several times in our acquisition. In these experiments, 1.0 × 10^8^ microbubbles (Micromarker® Fujifilm, Bracco) were injected in a bolus through a tail vein catheter.

For Fig. 4F-J, three xAM pulses were transmitted at sound sheet position 1 with a time between pulses set at 89 *μs*. These xAM pulses were next transmitted at sound sheet position 2. This sequence was repeated to acquire data for Doppler processing in two separate planes. A total of 2100 frames was acquired per sound sheet position with a frame rate of 1700Hz, leading to a 1.24s long acquisition. The average heart rate throughout the experiment was ranging from 300-360 bpm, leading to a cardiac cycle of 166-200 ms which is short enough to be covered in between 6-8 times during our acquisition. Here as well, 1.0 × 10^8^ microbubbles (Micromarker® Fujifilm, Bracco) were injected in a bolus through a vein catheter.

### Super-resolution ultrasound imaging

For the ULM and NSSLM experiments, similar surgical procedure and ultrasound pulse sequences were used than in the previous section. Ultrasound acquisitions were performed at a 1 kHz in two sound-sheet planes, recordings were continuous and lasted 105 seconds. Radiofrequency data for NSSLM was first filtered using the xAM operation and frames were beamformed using a delay-and-sum algorithm. For ULM, MB echoes were retieved using an SVD filter with a cutoff of 2 singular values and an additional high-pass filter was used to further demoise the data. For NSSLM, MB echoes were retrieved using a minimal SVD filter (cutoff of 1 singular value), and successive velocity filters. A radial symmetry localization algorithm was applied, followed by Kuhn Munkres pairing for trajectory reconstruction. For the composite rendering of NSSLM, two colorscales are applied based on different velocity domains.

## Acknowledgements

This project has been made possible in part by grants from the Delft University Fund, the 4TU Precision Medicine Program, the Medical Delta Ultra HB program, the Dutch Research Council (NWO.STU. 019.021 to DM), the Chan Zuckerberg Initiative (Dynamic RFA number 2023-321233 to DM) and the European Union (Marie-Sklodowska Curie Fellowship MIC-101032769 to BH). This research was also supported by the National Institutes of Health (grant R01-EB018975 to M.G.S.) and the Chan Zuckerberg Initiative. M.G.S. is a Howard Hughes Medical Institute Investigator.

