## Supplementary Materials for "Nonlinear sound-sheet microscopy: imaging opaque organs at the capillary and cellular scale"

### Figure 1: Experimental SSM and Orthogonal Plane Wave (OPW) linear imaging of a wire phantom

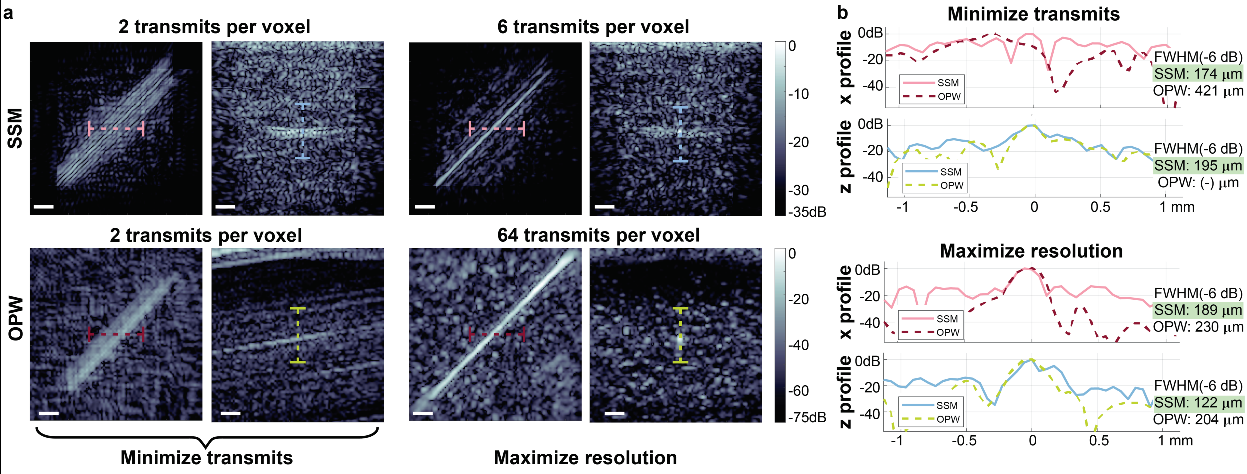


**a)** Experimental SSM and Orthogonal Plane Wave (OPW) linear imaging of a wire phantom. The first 2 columns are obtained by transmitting on each of the row and column array once (one single transmit per array). The top left image displays a C-scan obtained from multiple sound sheet micro scanning of the volume in both directions while the right panel displays the result on a single sound sheet. Images in the third and fourth column are obtained with three compounded angles for SSM and 2x32 angles for OPW and show C-scan and a single compounded sound sheet. **b)** Intensity profiles are plotted the SSM and OPW linear images with the color code indicated on overlay in the h panel.

### Figure 2: Imaging of wild type Anabaena and stripped Anabaena GVs

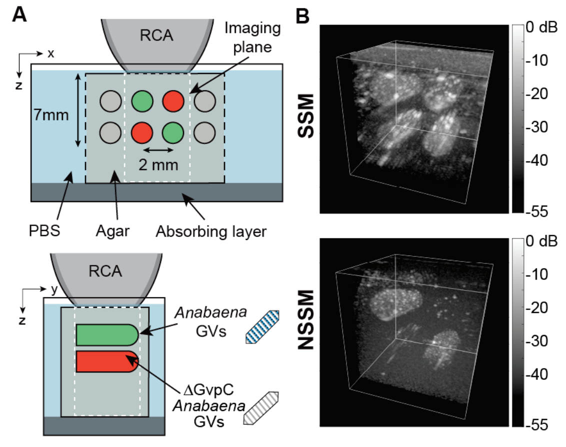


**a)** Schematic of the phantom configuration. An agar phantom (gray) immerged in phosphate buffered saline (PBS) contains wells filled with agar and linearly scattering *Anabaena flos aquae* gas vesicles and nonlinearly scattering *Anabaena flos aquae* gas vesicles stripped from the accessory protein GvpC . Top, cross-sectional view of the GV wells. Bottom, long axis view of the GV wells. **b)** $8.8x8.8x10mm^3$ volumes obtained from SSM (top) and NSSM (bottom) single direction micro scanning sequences (108 sound sheet positions). SSM clearly picks up increased signal in the wells compared to the surrounding agar while NSSM extinguishes linear signal successfully to retain the wells containing GVs without GvpC.

**Figure 3: (Nonlinear) Sound-Sheet Microscopy of perfused craniotomized rat brain**

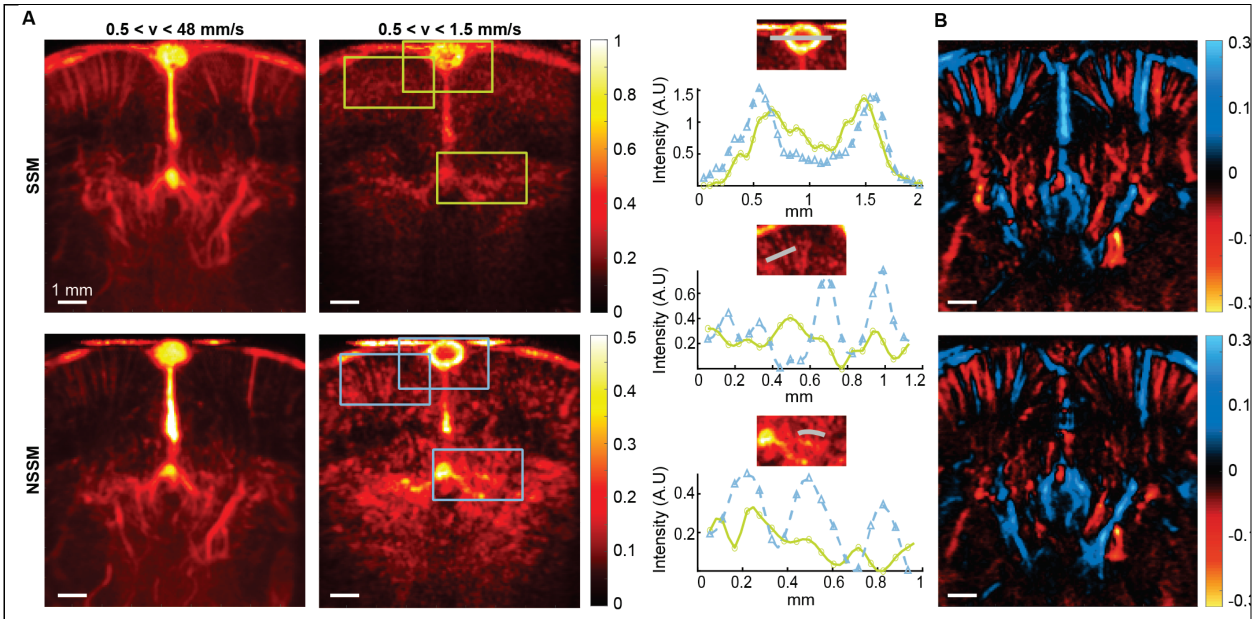


**a)** Frequency filtering of sound sheet Doppler frames for linear and nonlinear imaging sequences. The linear and nonlinear imaging sequence are obtained respectively before and after injection of microbubbles with a matched incident pressure. The left column shows axial Doppler velocity obtained after 10th order Butterworth band-pass filter with cutoff frequencies corresponding to velocity range [0.5-48] mm/s while the right column is focused on the range [0.5-1.5] mm/s pertaining to capillary flow. Scalebars are 1 mm. For the capillary flow, intensity profiles are taken at specific locations indicated by gray lines in the zoomed-in inserts on the right. The profiles in green and blue show intensities (in arbitrary units) taken respectively in the linear and nonlinear sequence. **b)** Signed axial Doppler velocity obtained from linear (top) and nonlinear sound sheet sequence. The positive velocities (indicated in blue) are taken as upwards velocity (towards the probe). In the cortical areas, this color code shows distinction between veins (blue) and arteries (red).

**Figure 4: Delays and apodizations used for high-speed multi-view Doppler NSSM**

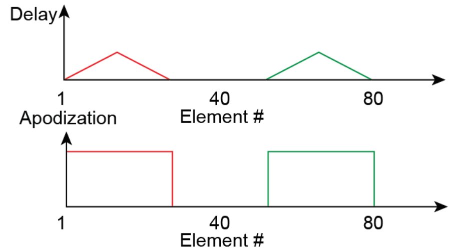


Ultrasound insonification scheme for multi slice NSSM in consecutive cardiac cycles. Two sound sheet positions can be arbitrarily set at positions as far as $d = 6\ mm$, with a theoretical pulse repetition frequency of $37\ kHz$ that enables ultrafast Doppler imaging in two sound sheet planes, with a maximum framerate attainable of close to 4200 Hz. In this setup, we limit the framerate to 1700 Hz. The delays and apodization of each sound sheet are represented for an angle of 18 degrees. For the

NSSI mode, the 3 pulses necessary for amplitude modulation are fired for one sound sheet, and then the 3 pulses for the next sound sheet are fired. This sequence is then looped over the number of frames.

**Figure 5: High-speed multi-view NSSM spectrograms of the rat brain vasculature**

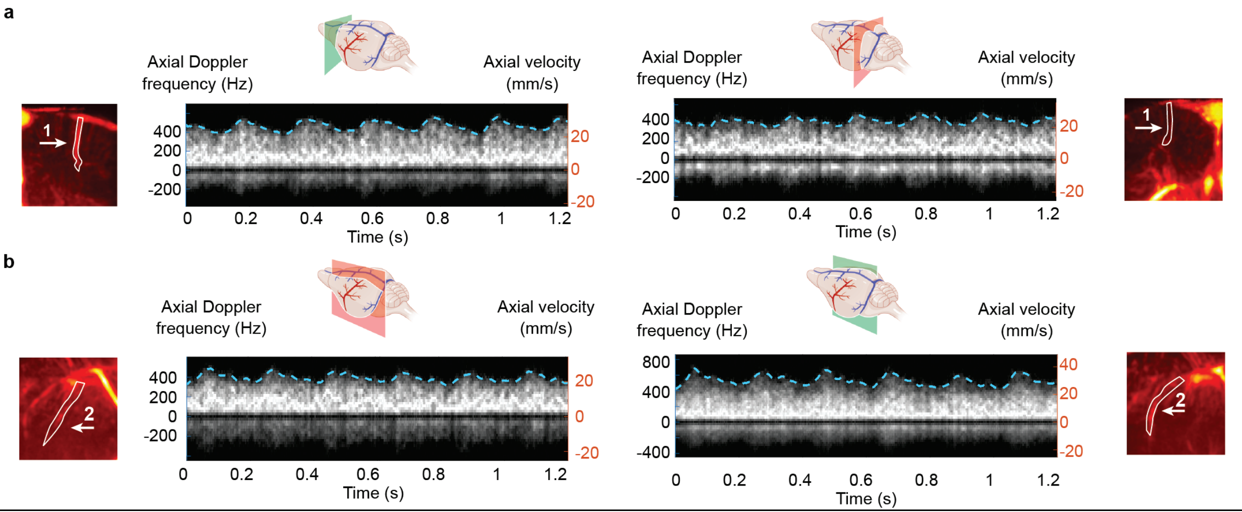


**a)** Doppler spectrum revealing peak cerebral blood flow and pulsatility of a cortical arteriole of the right hemisphere in the first (green) and second (red) plane. **b)** Doppler spectrum revealing peak cerebral blood flow and pulsatility of a sub-cortical vessel of the left (red) and right (green) hemisphere.

**Figure 6: NSSLM composite rendering for the second sound-sheet of the multi-view acquisition in the coronal plane**

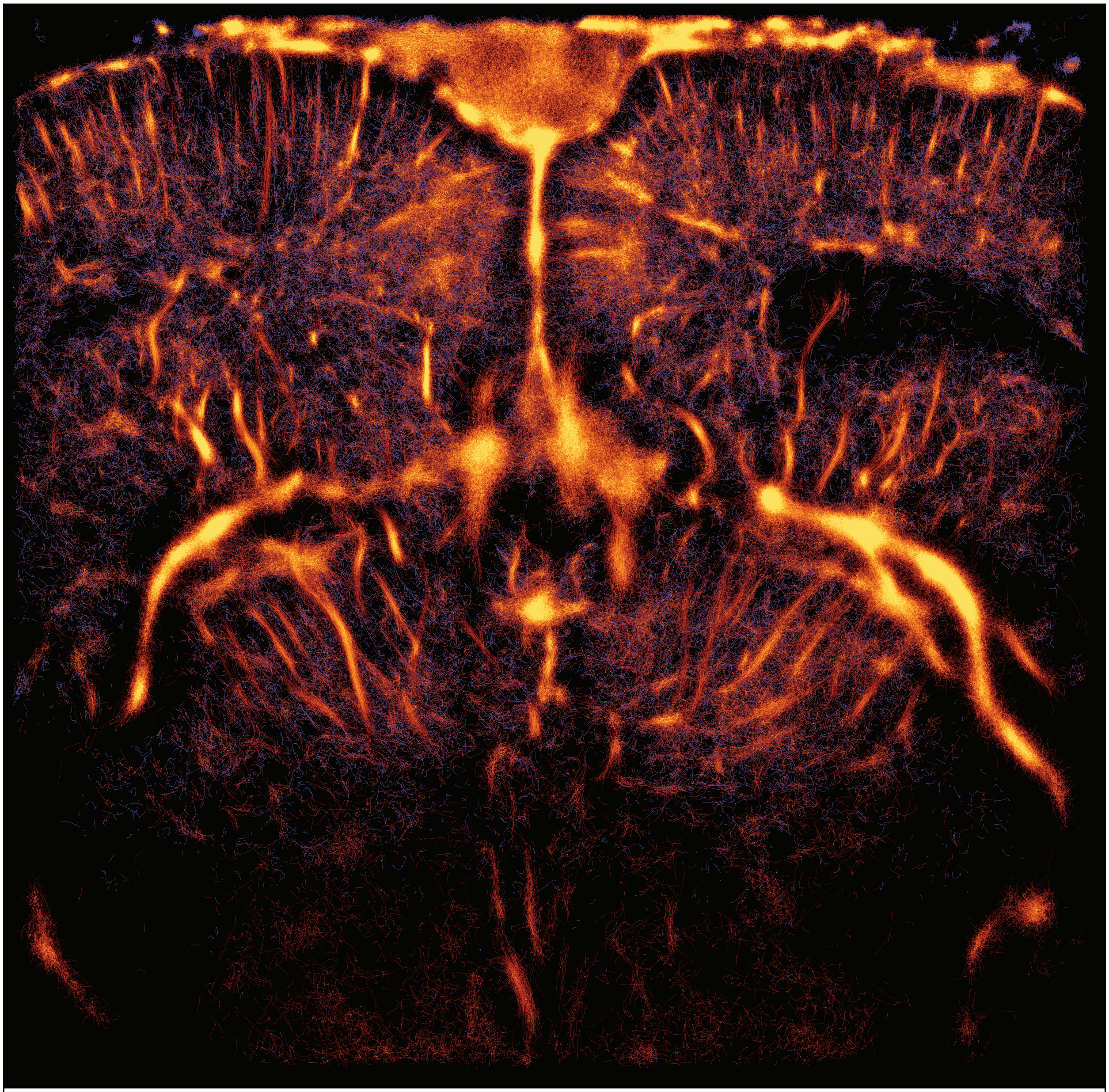

Composite NSSLM image display all cerebral blood flow velocity bands for the second multi-view plane

**Figure 7: SSLM rendering for the second sound-sheet of the multi-view acquisition in the coronal plane**

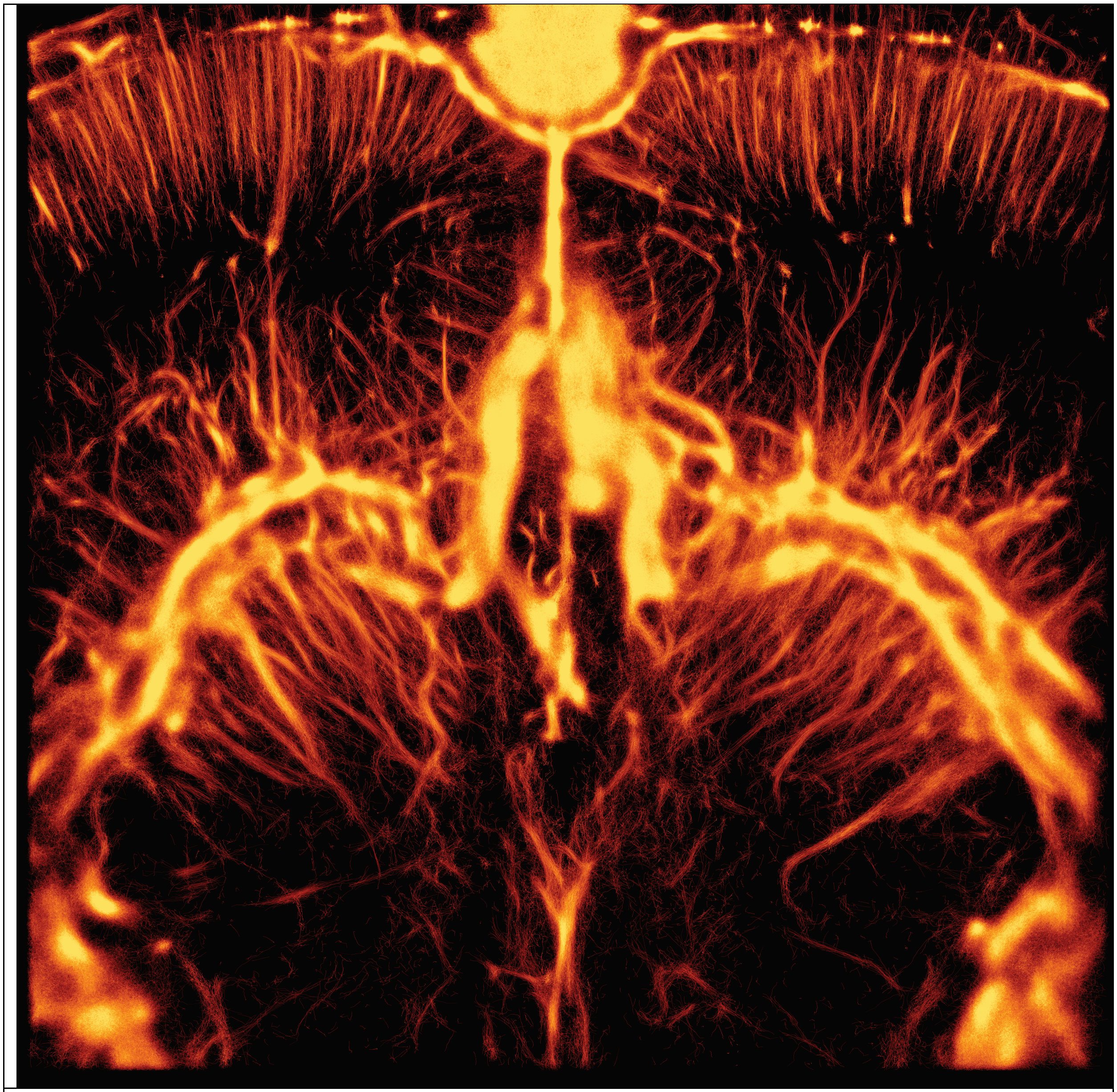

SSLM image generated with a state-of-the-art ULM processing pipeline.
